# Modelling functional human neuromuscular junctions in a differentially-perturbable microfluidic environment, validated through recombinant monosynaptic pseudotyped ΔG-rabies virus tracing

**DOI:** 10.1101/745513

**Authors:** Ulrich Stefan Bauer, Rosanne van de Wijdeven, Rajeevkumar Nair Raveendran, Vegard Fiskum, Clifford Kentros, Ioanna Sandvig, Axel Sandvig

## Abstract

Compartmentalized microfluidic culture systems provide new perspectives in *in vitro* disease modelling as they enable co-culture of different relevant cell types in interconnected but fluidically isolated microenvironments. Such systems are thus particularly interesting in the context of *in vitro* modelling of mechanistic aspects of neurodegenerative diseases such as amyotrophic lateral sclerosis, which progressively affect the function of neuromuscular junctions, as they enable the co-culture of motor neurons and muscle cells in separate, but interconnected compartments. In combination with cell reprogramming technologies for the generation of human (including patient-specific) motor neurons, microfluidic platforms can thus become important research tools in preclinical studies. In this study, we present the application of a microfluidic chip with a differentially-perturbable microenvironment as a platform for establishing functional neuromuscular junctions using human induced pluripotent stem cell derived motor neurons and human myotubes. As a novel approach, we demonstrate the functionality of the platform using a designer pseudotyped ΔG-rabies virus for retrograde monosynaptic tracing.

**Graphical abstract:** 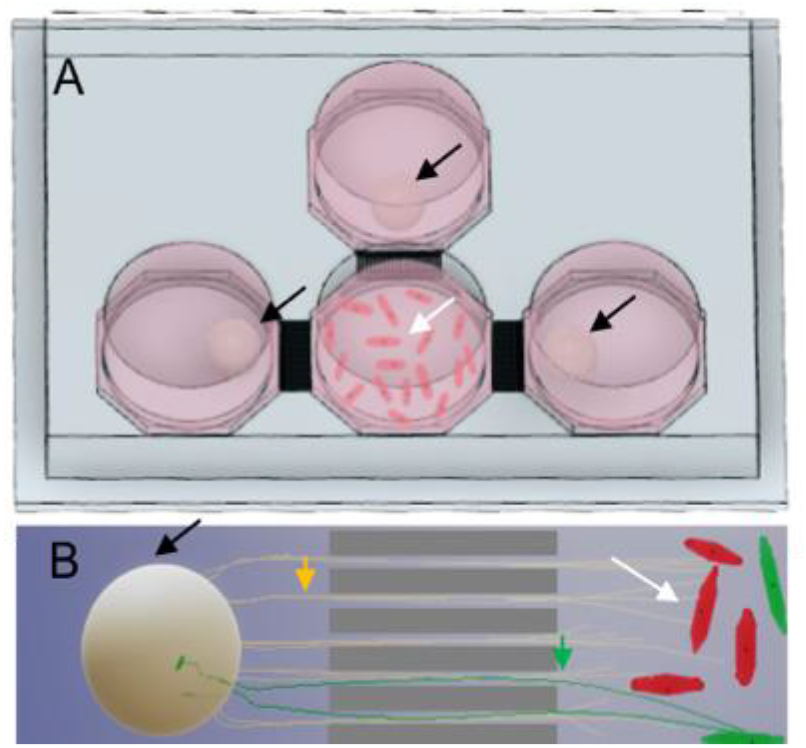
Functional neuromuscular junction in a microfluidic chip. **(a) Overview of microfluidic chip**. Human iPS cell-derived motor neuron aggregates (spheroids indicated by black arrows) are seeded in the three lateral compartments of the chip, while human myotubes (white arrows) are seeded in the middle compartment. **(b) Directed connectivity and retrograde virus tracing**. Outgrowing axons (yellow arrow) from the motor neuron aggregate enter the directional axon tunnels (grey rectangles) and form connections with the myotubes (white arrow) within the opposite compartment. Addition of a designer monosynaptic pseudotyped ΔG-rabies virus to the myotube compartment, infects the myotubes (green) expressing an exogenous receptor (TVA) and rabies glycoprotein (G), subsequently making infectious viruses that are retrogradely transported through the motor neuron axons (green arrow) back to the neuronal cell bodies within the aggregate, validating neuromuscular junction functionality.

## Introduction

Amyotrophic Lateral Sclerosis (ALS) is a progressive, fatal neurodegenerative disease that primarily affects upper and lower motor neurons (MN) with a lifetime risk of 1:350 in men and 1:400 in women, making it the third most prevalent neurodegenerative disease.^1,2^ The progressive degeneration and, ultimately, loss of motor neurons affects synaptic transmission at the neuromuscular junction (NMJ). As a result, early symptoms of the disease can manifest as muscle twitches or cramps and gradually progress to include spasticity, leading to muscle atrophy, loss of control of voluntary movements, and inevitably respiratory failure, typically within 3-5 years after symptom onset.^3,4^ A wide range of mutations have been identified as causing the familial form of the disease, but while our knowledge concerning underlying mutations has been rapidly expanding in recent years, our understanding of the molecular pathology, particularly the onset and mechanism by which the respective mutations cause the disease, remains lackluster. Similarly, the mechanisms underlying the sporadic form of the disease are poorly understood.

Current preclinical *in vivo* models of ALS make use of transgenic mice overexpressing some of the most common gene mutations associated with ALS, typically SOD1, TDP-43, FUS, or C9ORF72. Each model has distinctive advantages, however, no single model faithfully mimics disease complexity (Nardo et al, 2016; Kabashi et al, 2008; Sharma et al, 2016; Philips and Rothstein, 2016; O’Rourke et al, 2015).^3,5–8^ An alternative or complementary approach to *in vivo* ALS models, is presented by *in vitro* cell-based models. In the last decade, such *in vitro* models have undergone significant development, especially after the advent of cell reprogramming technologies enabling the conversion of somatic cells into induced pluripotent stem (iPS) cells after transduction with only four genes (Oct4, Sox2, Klf4 and c-Myc).^9^ Furthermore, iPSCs can be differentiated into specific neuronal subtypes, including MNs.^10–12^

Recent developments in cell culture platforms, such as microelectrode arrays (MEAs) and microfluidic chips provide new options for investigating the electrophysiological behaviour of neural networks.^13,14^ In the context of ALS, of particular relevance are models based on the recapitulation of the NMJ in healthy and perturbed conditions. Indeed, a number of studies have demonstrated the applicability of various bioengineering methods for creating muscle-neuron co-cultures, including functional NMJs *in vitro*.^15–21^ In the present study, we model functional human NMJs by compartmentalized culture of iPS cell-derived MN pseudo-organoids and myotubes in a differentially perturbable multi-nodal microfluidic chip and demonstrate that NMJ function can be validated through monosynaptic retrograde tracing using a designer pseudotyped ΔG-rabies virus (RV).

## Materials and Methods

Induced pluripotent stem (iPS) cells were purchased from Takara Bio Europe AB (Y00305). The cells were expanded on Laminin 521 (LN521, BioLamina) coated flasks using iPS expansion media containing Knockout-DMEM/F-12 (12660012, Invitrogen), 20% Xeno-free CTS Knockout Serum Replacement (A1099202, ThermoFisher), Penicillin-streptomycin, Non-essential amino acids, L-glutamine, 2-Mercaptoethanol 55mM (21985-023, ThermoFisher), and 4ng/ml FGF-Basic (AA 10-155) Recombinant Human Protein (PHG0026, Life Technologies).

A modified version of the protocol published by Amoroso et al.^12^ was used to differentiate iPS cells to motor neurons. iPSC-derived motor neurons were maintained in motor neuron media (MN media) consisting of Neurobasal Medium (21103049, ThermoFisher), N2 supplement (17502-048, Gibco), B27 supplement (17504-044, Gibco), Non-essential amino acids, L-glutamine, Penicillin-streptomycin, 0.4μg/ml Ascorbic acid, 25μM L-glutamate, 1μM all-trans Retinoic acid, 25μM 2-Mercaptoethanol, 20μM Y-27632 (Y0503, Sigma-Aldrich).

Primary human skeletal myoblasts (A11440, ThermoFisher) were expanded in DMEM +L-glutamine (Gibco,21041-025), 20% Embryonic stem-cell FBS (Gibco, 16141061), Dexamethasone, Insulin (human recombinant expressed in yeast, (I-2643, Sigma-Aldrich), 20 ng/ml EGF Recombinant Human Protein (10605HNAE250, Life Technologies), 20ng/ml FGF-Basic (AA 10-155) Recombinant Human Protein, 1:100 Penicillin-streptomycin.

Myoblast to myotube differentiation was achieved by switching to myoblast differentiation media (MDM) DMEM basal medium (11885-084, ThermoFisher), 2% Horse Serum (16050-130, ThermoFisher), Penicillin-streptomycin.

### iPS cell-derived motor neuron (MN) cultures on MEAs

Five 60EcoMEA-Glass microelectrode arrays (Multichannel systems) were disinfected with 70% ethanol for 5 minutes or less and washed with PBS before sterilizing under ultraviolet light overnight. The MEAs were then treated with foetal bovine serum (FBS) overnight to ensure the surfaces were hydrophilic. After removing the FBS, the MEAs were coated with Matrigel (Sigma-Aldrich), diluted 1:30 in cell expansion media. Each MEA was coated with 750µL of diluted Matrigel and left for 30 minutes at 4°C, followed by 30 minutes at room temperature. After removing the coating, the MEAs were seeded with 200µL of rat astrocytes at a concentration of 100 000 cells/mL, which means each MEA was seeded with approximately 20 000 astrocytes. 3 days after seeding the astrocytes, approximately 100 000 MNs were seeded onto each MEA at a density of 1 750 000 cells/mL. Culture media was replaced 3 times per week up until culture age of 28 days, after which media was replaced once per week.

### *In vitro* electrophysiology and antigenic profile of MN networks

Recordings of electrical activity of the motor neurons were recorded on day 8, 15, 22, 28, 35, 42, 49, 56, 63 and 70 days *in vitro* (DIV). Each recording was 4 minutes long at 10 000Hz, and was made at least 24 hours after media replacement. Only baseline activity was recorded with no stimulation. Five cultures were maintained up until 42DIV, but only four of these could be maintained up until 70 DIV. Spike detection was performed in NeuroExplorer 5, after filtering the recorded signal through a fourth order Butterworth filter from 300 to 3000Hz, and spikes were identified in signals that were greater than four standard deviations away from the mean signal, and the timestamps of spikes were recorded. These spike trains were exported to MatLab R2018a (MathWorks) for further analysis.

In order to confirm the identity of the cells as MNs, cells were cultured in parallel to the MEAs in 4 wells on an 8-well Ibidi chip. At 53DIV, these were stained for motor neuron markers. First, the cell media was aspirated, and the cells were washed with warm PBS. Then they were fixed with 4% paraformaldehyde for 15-25 minutes before being treated with a block solution of 5% goat serum and 0.3% Triton-X in PBS for 1-2 hours. After removing the block, three wells were stained with primary antibodies for Islet 1 (Abcam, ab86472, 1:250 dilution), HB9 (Abcam, ab221884, 1:2500 dilution) and heavy neurofilament (Abcam, ab4680, 1:1000 dilution) with 1% goat serum and 0.1% Triton-X in PBS, while the last well was used as a control, filled only with PBS with goat serum and Triton-X. The primary antibody was left overnight at 4°C. The next day, the wells were washed with PBS 3 times 10 minutes, before adding secondary antibodies (ThermoFisher, A11001, A11036, A21449, all at 1:1000 dilution) with 1% goat serum and 0.1% Triton-X in PBS to 3 out of the 4 wells, including to the well which was not stained with the primary antibodies. The last well was filled with only PBS with goat serum and Triton-X, so that there was a control with primary antibody only and one with secondary antibody only. The wells were covered in aluminium foil to prevent light exposure and left for 2 hours, and Hoechst staining added (Sigma-Aldrich 14533, 20µg/mL at 1:10 000 dilution) during the last 5 minutes. After removing the secondary antibodies, the wells were washed 3 times 15 minutes before covering the cells with a glass coverslip using Fluoroshield Mounting Medium (Abcam ab104135). The finished slides were then left at 4°C overnight. When imaging the cells, exposure time was set to 300-400ms for all stains except Hoechst, which was set to 8-15ms.

### Microfluidic chip design and production

In this work, a semi-open system microfluidic chip is used that is comprised of four cell compartments interconnected through micro-sized tunnels only permissible to axons. In a recent publication, we demonstrated the compatibility of such a microfluidic chip with neuronal aggregates (van de Wijdeven et al. 2018). A photoresist mould was fabricated using standard lithographic techniques, more details regarding the design and the fabrication procedure can be found in a previous study (van de Wijdeven et al. 2018). Poly(dimethylsiloxane) (PDMS) was cast on top of the mould and cured in an over at 65 °C for 4 hours. Afterward, the PDMS was carefully peeled from the mould and the cell compartments were opened using a punching device (Ø 6mm). The PDMS debris was removed with adhesive tape and consecutive washes in acetone, ethanol (70%) and deionized DI water. The chips were semi-dried with an airgun and left to dry overnight. Subsequently, the chips were irreversibly bonded onto glass coverslips (24 × 32 mm Menzel-Gläser) by exposing both surfaces to oxygen plasma for 1 min followed by directly heating the assembled device at 75 °C for 30 seconds. Finally, the chips were filled with DI water and kept at 4 °C for storage. Prior to coating, the chips were sterilized under UV light.

### Immunocytochemistry of *in vitro* neuromuscular junctions

Cells were fixed in 4% Parafomaldehyde in PBS, blocking was carried out with 5% Goat serum and 0.6% Triton-X in PBS and primary and secondary immunostaining in 2.5% Goat serum and 0.3% Triton-X in PBS (for first BTX staining and staining after the non-blinded BTX activity abolishing experiment 0.06% and 0.03% Triton-X were used). Immunostainings were carried out using 1:50 mouse anti-Synaptophysin (ab8049, abcam), 1:100 rabbit anti-Troponin I (701585, ThermoFisher), 1:200 mouse anti-Troponin T (SAB420717, Sigma-Aldrich), 1:5000 chicken Neurofilament-Heavy (ab4680, abcam), 1:100 rabbit anti-Nicotinic Acetylcholine Receptor alpha 1 (ab221868, abcam), 1:2500 chicken anti-Beta III Tubulin (ab41489, abcam), 1:50 rabbit anti-HB9 (ab221884, abcam), 1:50 rabbit anti-Islet1 (ab109517, abcam), 1:1000 mouse anti-NeuN (ab104224, abcam), 1:20 rabbit anti-Galactocerebroside (AB142, Sigma-Aldrich), 1:1000 mouse anti-Beta III Tubulin (ab119100, abcam), and 1:5000 chicken anti-Glial fibrillary acidic protein (ab4674, abcam). Nuclear staining was carried out using Hoechst DNA stain (bisBenzimide H 33342 trihydrochloride, 14533, Sigma-Aldrich)

Secondary antibody staining was carried out using 1:500 Alexa Fluor 488 goat anti-mouse IgG antibody (A-11001, Life Technologies), 1:500 Alexa Fluor 488 goat anti-rabbit (A-11008, Life Technologies), 1:500 Alexa Fluor 568 goat anti-mouse IgG antibody (A-11019, Life Technologies), 1:500 Alexa Fluor 568 goat anti-rabbit IgG antibody (A-11079, Life Technologies), 1:500 Alexa Fluor 647 goat anti-rabbit (A-21244, Life Technologies), and 1:500 Alexa Fluor 647 goat anti-chicken (A-21449, Life Technologies).

Alpha-Bungarotoxin (α-BTX) labelling was carried out simultaneously with secondary antibody labelling. A 1mg/ml stock of Alexa Fluor™ 488 conjugate of α-Bungarotoxin (B13422, ThermoFisher) was used at 1:100 dilution in the secondary antibody solution.

### Alpha-Bungarotoxin (α-BTX) functional assay

Chips seeded as described above were allowed to mature for 21 days after seeding. Chips were selected for high contractile activity of myotubes that had visual contact with axons (more than 1 contraction per minute over 5 minutes with at least 1 contraction in each minute, the day prior to the experiment). Video of pre-experiment activity was recorded for 1 min followed by counting of the total number of contractions in 10 min. The cells were allowed to recover for 1 h at 37°C, 5% CO_2_ before addition of 1:100 of α-BTX at a final concentration of 1.25μM or of 0.75μl sterile PBS and incubation of 10 min at 37°C, 5% CO_2_. The α-BTX was only added to the central well containing the myotubes, which was fluidically isolated by hydrostatic pressure throughout the incubation. Another 1 min video of activity after intervention was recorded before counting the total number of contractions in 10 min.

Cells were fixed immediately after the final count. For quantitative analysis the experiment was repeated with 10 min video being recorded as baseline followed by 1 h recovery at 37°C, 5% CO_2_ and treatment. Another 10 min video was recorded after treatment, and blinded quantification was carried out for both.

### EnvA pseudotyped-G deleted rabies virus ((ΔG-RV) production

Endotoxin free plasmid maxipreps (Qiagen) were made for all transfections. pAAV-CMV-TVAmCherry-2A-oG (#104330) and pCAG-YTB (#26721) for expressing TVA and Rabies Glycoprotein were purchased from Addgene.

EnvA pseudotyped-G deleted rabies virus expressing GFP (EnvA-ΔG-RV-GFP) or mCherry (EnvA-ΔG-RV-mCherry) transgene were produced as described in Wickersham et al Nat Protocols 2010^22^, Osakada and Callaway Nat Protocols 2013^23^. Briefly, for the recovery of ΔG-RV, 293T cells were cotransfected with the rabies genome construct cSPBN-4GFP (Addgene #52487) along with helper plasmids pcDNA-SADB19N (#32630), pcDNA-SADB19P (#32631), pcDNASADB19L (#32632), pcDNA-SADB19G (#32633) and pCAGGS T7 (#59926). Transfection was carried out using Lipofectamine 2000 (ThermoFisher). The transfected 293T cells were maintained in 10% FBS/DMEM in a humidified atmosphere of 5% CO2 at 37oC. ΔG-RV particles rescued from the cDNA were amplified by passaging for several days on a complementing cell line BHK-B19G2 cells. For pseudotyping with EnvA, BHK-EnvA cells were infected with unpseudotyped ΔG-RV at a multiplicity of infection (MOI) of 0.5. Twenty-four hours post-infection, infected cells were washed with DPBS, trypsinized with 0.25% trypsin-EDTA and reseeded on multiple 15 cm dishes in a humidified atmosphere of 3% CO2 at 35oC as described. Incubation medium was harvested 48 hrs later, filtered using 0.45 µm sterile filter and viral particles were concentrated by ultracentrifugation at approximately 50,000g for 2 hrs at 4oC, for producing high titer EnvA-pseudotyped ΔG-RV. The pellets were resuspended in DPBS and concentrated using Amicon Ultra centrifugal filters (Millipore). Unpseudotyped ΔG-RV and EnvA-pseudotyped ΔG-RV were titrated by serial dilutions using HEK293T cells and HEK293-TVA800 cells, respectively. Titer of viral stocks EnvA-ΔG-RV-GFP and EnvA-ΔG-RV-mCherry were determined as approximately 1010 infectious particles/ml.

The pseudotyped ΔG-rabies virus was produced by the Viral Vector Core Facility, Kavli Institute for Systems Neurosceince, NTNU, Norway.

### Retrograde tracing with RV *in vitro*

The same overall layout in the microfluidic chips as in previous experiments was used. The myoblasts were expanded and nucleofected at P2 using P5 Primary Cell 4D-Nucleofector^®^ X Kit L (Lonza, V4XP-5012) with the pCAG-YTB (26721, Addgene) or pAAV-CMV-TVAmCherry-2A-oG (104330, Addgene) that expresses exogenous receptor TVA and the rabies glycoprotein (B19G or optimized G) or pmaxGFP as control (part of the P5 Primary Cell 4D-Nucleofector^®^ X Kit L, V4XP-5024, Lonza) plasmids using the EY-100 setting in the Lonza 4D-Nucleofector™ Core Unit (AAF-1002B, Lonza) and 4D-Nucleofector™ X Unit (AAF-1002X, Lonza). Cultures were allowed to develop without interference until 15 days in vitro (DIV) with full media changes every other day. At 15 DIV, the pAAV-CMV-TVAmCherry-2A-oG-nucleofected or pCAG-YTB-nucleofected myotubes were respectively infected with EnvA-ΔG-RV-GFP or EnvA-ΔG-RV-mCherry virus in myotube differentiation media overnight. After incubation, the media were removed and the wells were washed once with warm MDM and again filled with MDM. Full media changes were carried out every other day thereafter.

pAAV-CMV-TVAmCherry-2A-oG was a gift from Marco Tripodi (Addgene plasmid #104330; http://n2t.net/addgene:104330; RRID:Addgene_104330), pCAG-YTB was a gift from Edward Callaway (Addgene plasmid #26721; http://n2t.net/addgene:26721; RRID:Addgene_26721).

## Results

### Spontaneous electrical activity profile of iPS cell-derived MN networks on MEAs

Analysis of the data obtained from electrophysiological recordings of iPS cell-derived MN networks on 60-electrode MEAs revealed the spontaneous electrical activity profile of these networks over time. Based on the firing rate of these MNs across the entire MEA and their burstiness index (BI), we could classify the activity of the MNs into three categories: (i) “Young” networks, 8-35 DIV characterised by low, but increasing, firing rate and low BI; (ii) “Middle aged” networks, 42-56 DIV showing high firing rate and medium BI; and (iii) “old” networks showing low firing rate and high BI (Figure 1).

**Figure 1.**
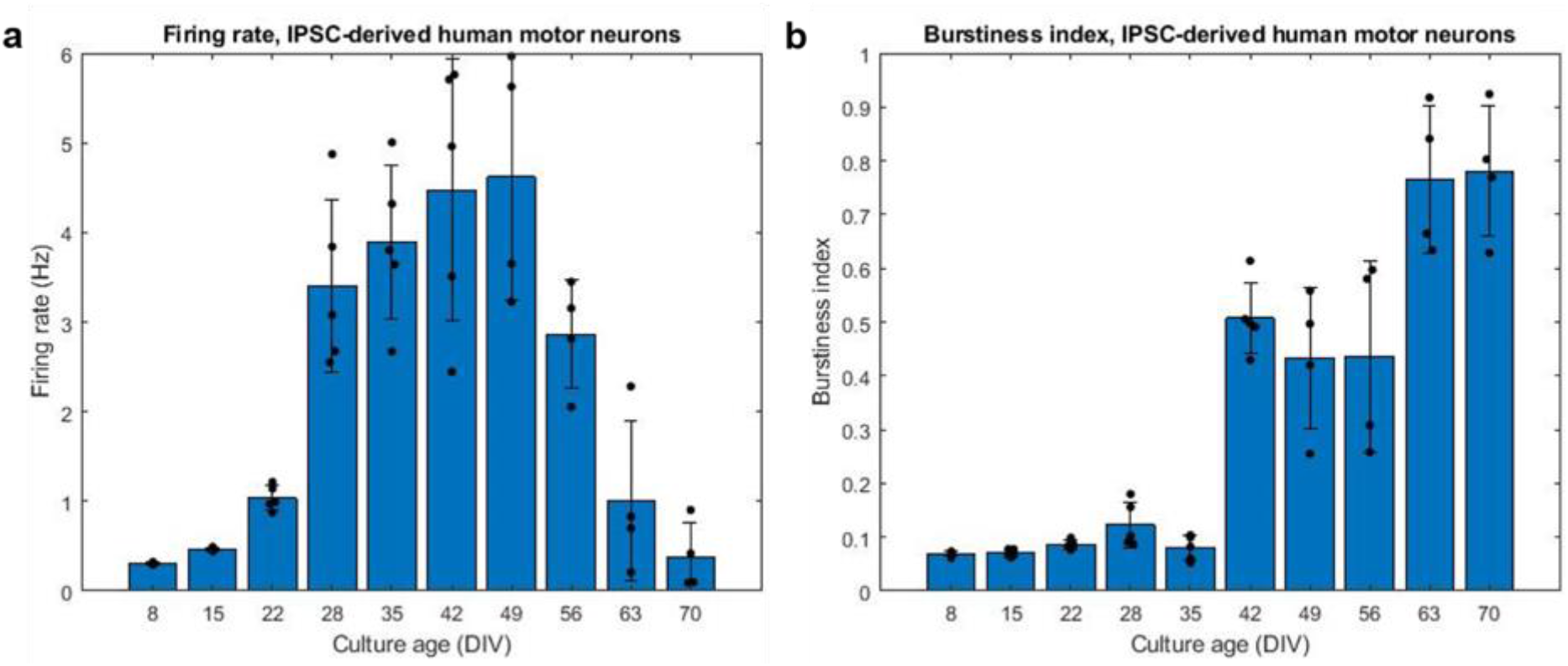
Development of MN network activity. Development of activity of networks of IPSC-derived motor neurons. (**a**) Firing rate increased until 49DIV, at which point it decreased quickly. (**b**) Burstiness index increased for as long as the cultures were maintained. N=5 for culture age 8-42DIV, N=4 from 49-70DIV.

Immunocytochemistry of iPS cell-derived MNs clearly showed the presence of MN-specific markers Islet 1 and Homeobox gene HB9, as well as the neuron specific marker neurofilament heavy (NEFH). Figure 2 shows the expression of these proteins at two different scales, clearly demonstrating presence and overlap of Islet 1 and HB9 together with NEFH, providing clear evidence of the presence of MNs.

**Figure 2.**
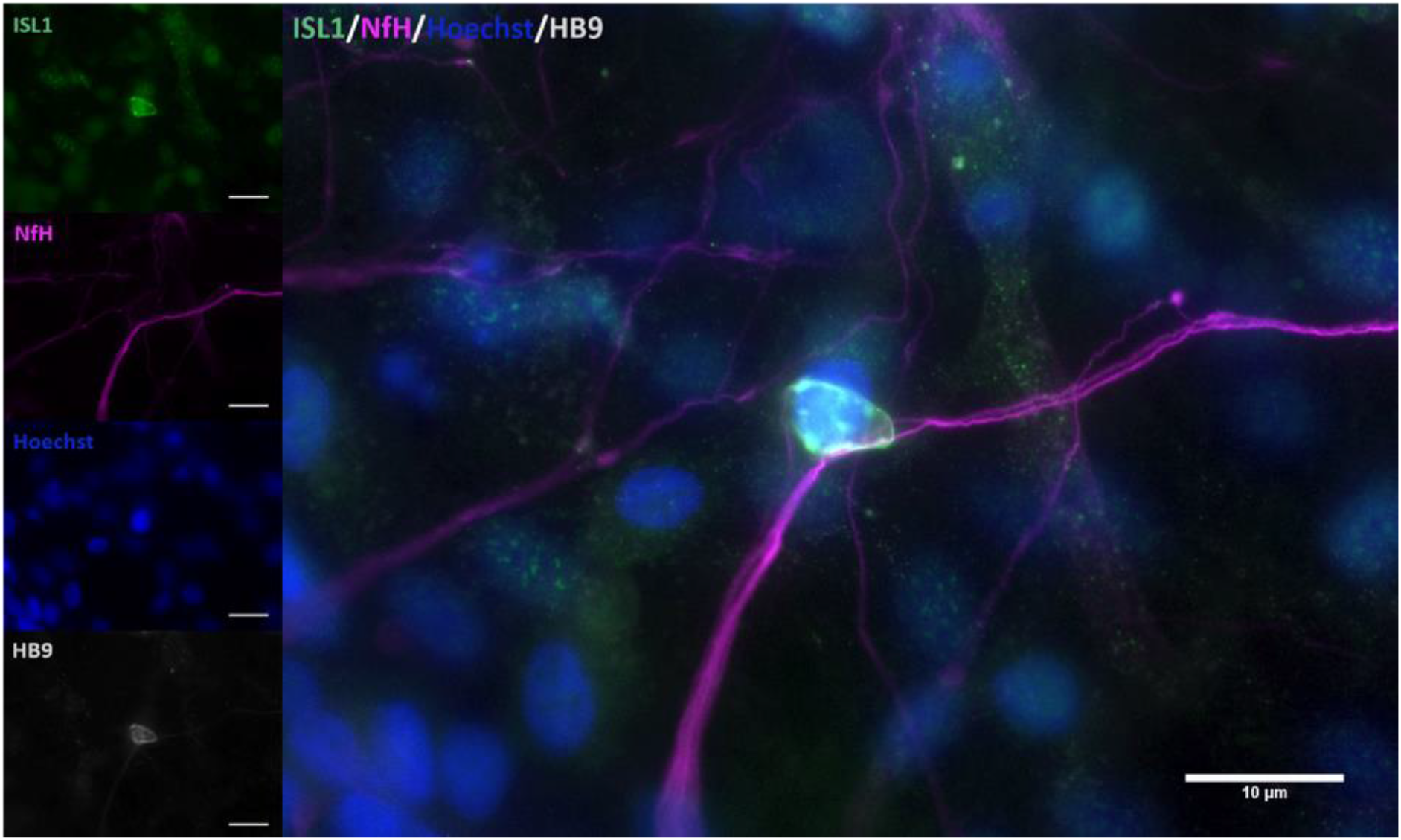
Antigenic profile of iPS cell-derived MNs. Staining for MN specific markers clearly verified their presence in the cell cultures. Clear overlap was seen between Islet 1 (green) and HB9 (grey), confirming the identity of the stained cells. Staining for Hoechst (blue) also showed that cells which did not stain positive for MN markers were also present in the cultures.

We cannot exclude the presence of other cell types, given the large number of cells identified by Hoechst that did not show any expression of neuronal markers. Primary or secondary antibody exclusion assays showed no immunostaining (data not shown).

### NMJ formation within microfluidic chips

To establish *in vitro* NMJs, pseudo-organoids of the human iPS cell-derived MNs (Figure 3) and human myotubes were seeded in separate cell compartments in an in-house developed multi-nodal microfluidic chip with directional tunnels connecting the compartment housing the pseudo-organoids with the one containing the myotubes.^13^ MN axon outgrowth began promptly and, typically within 3 days in chips (DIC), the first axons had reached through the tunnels into the adjacent cell compartment (Figure 4). By 8 DIC the first contracting myotubes could be observed. Larger myotubes were also observed developing spontaneous activity without direct contact with MN axons. The diameter and length of the axon bundles exiting the tunnels and the number of myotubes contacted by these axons continued to increase until all compartments were fixed after 21 DIC. Although some cells from the pseudo-organoids settled onto the bottom of the well and migrated away, the vast majority of cells remained within the pseudo-organoids.

**Figure 3.**
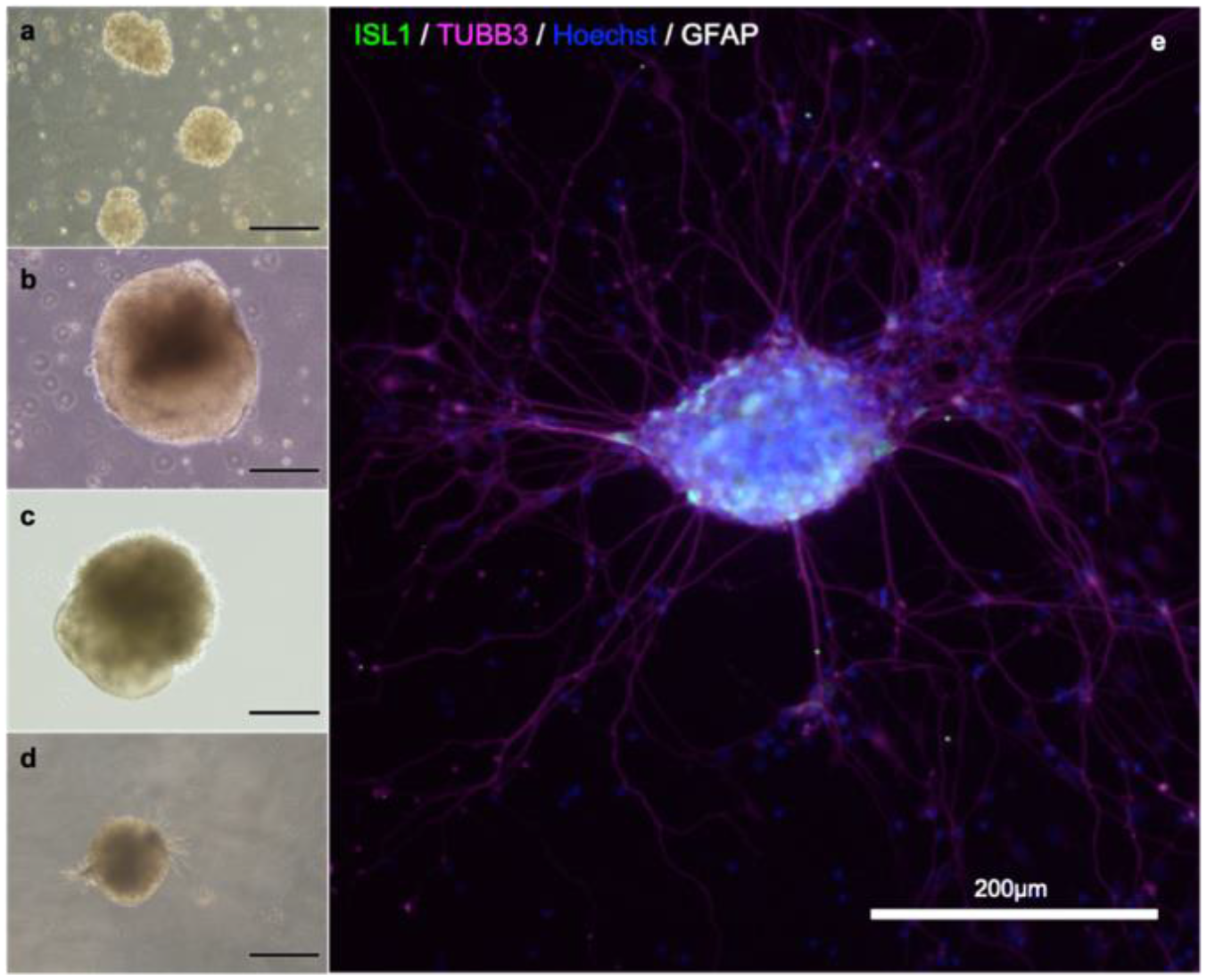
MN aggregate derived from iPS cell reprogramming. Cells during differentiation, corresponding to day 2 (**a**), 9 (**b**), 18 (**c**) and 25 (**d**). After 21 days, MNs are present in the population. Neuronal aggregates (**e**) were formed after dissociation at day 30 and post-seeding and maturation for another 9 days. Immunostaining of the aggregates reveals the presence of immature neurons (β-III-tubulin; magenta), motor neurons (Islet 1; green), and also some astrocytes (GFAP; grey). Nuclear counterstaining with Hoechst (blue).

**Figure 4.**
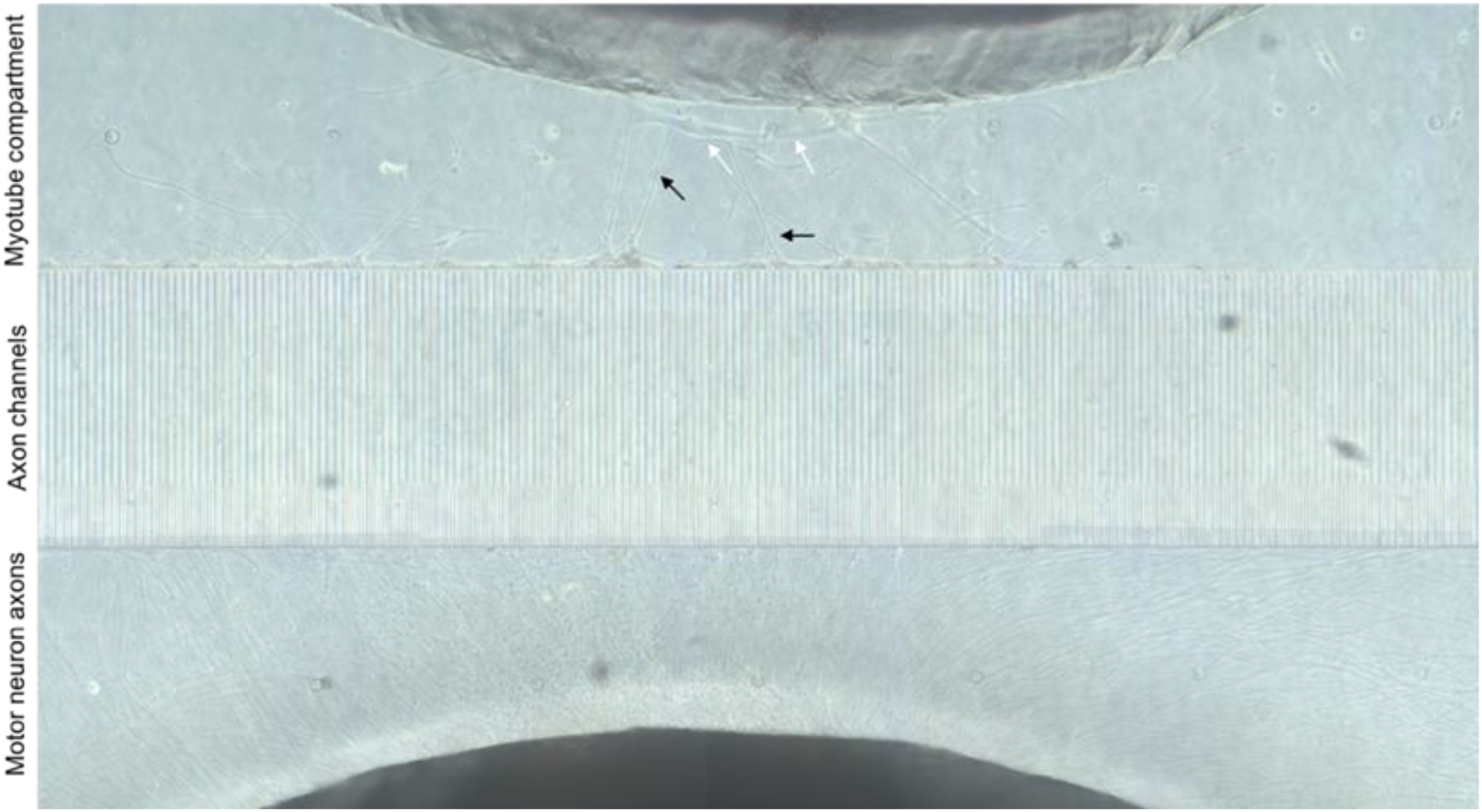
Directional MN axon outgrowth into the muotube compartment. Representative image from the microfluidic chips at 8 DIV, showing directional axonal outgrowth from the MN aggregate into the axon tunnels and entry into the opposite compartment containing myotubes. MN neuron axons (black arrows) are clearly visible and appear to contact myotubes (white arrows).

### Verification of synaptic contact between MNs and myotubes

Immunostaining within the microfluidic chips confirmed expression of synaptophysin (SYP), Troponin I (TnI), Troponin T (TnT), Neurofilament heavy (NFH), and beta-III-tubulin (Figure 4). Synaptophysin and CHRNA3 (cholinergic receptor nicotinic alpha 3 subunit) expression could be observed around the surface of the troponin-positive cells (Figure 5), suggesting the presence of NMJs. Additional immunostaining combined with staining using a fluorescent conjugate of the postsynaptic nicotinic acetylcholine receptor-specific Bungarus multicinctus toxin-derived long-chain α-neurotoxin α-Bungarotoxin (BTX) revealed BTX expression specific to the surface of myotubes that were in contact with axons, as well as expression of skeletal muscle proteins TnI or TnT (Figure 6). Furthermore, most of the myotubes with BTX staining on their surface had been observed contracting before the cultures were fixed. In addition to the above, the presence of mature MNs in the pseudo-organoids was further verified with immunocytochemistry of pseudo-organoid sections revealing cells co-expressing NeuN and HB9 (Supplementary figure 1) as well as NeuN and Islet1 (Supplementary figure 1).

**Figure 5.**
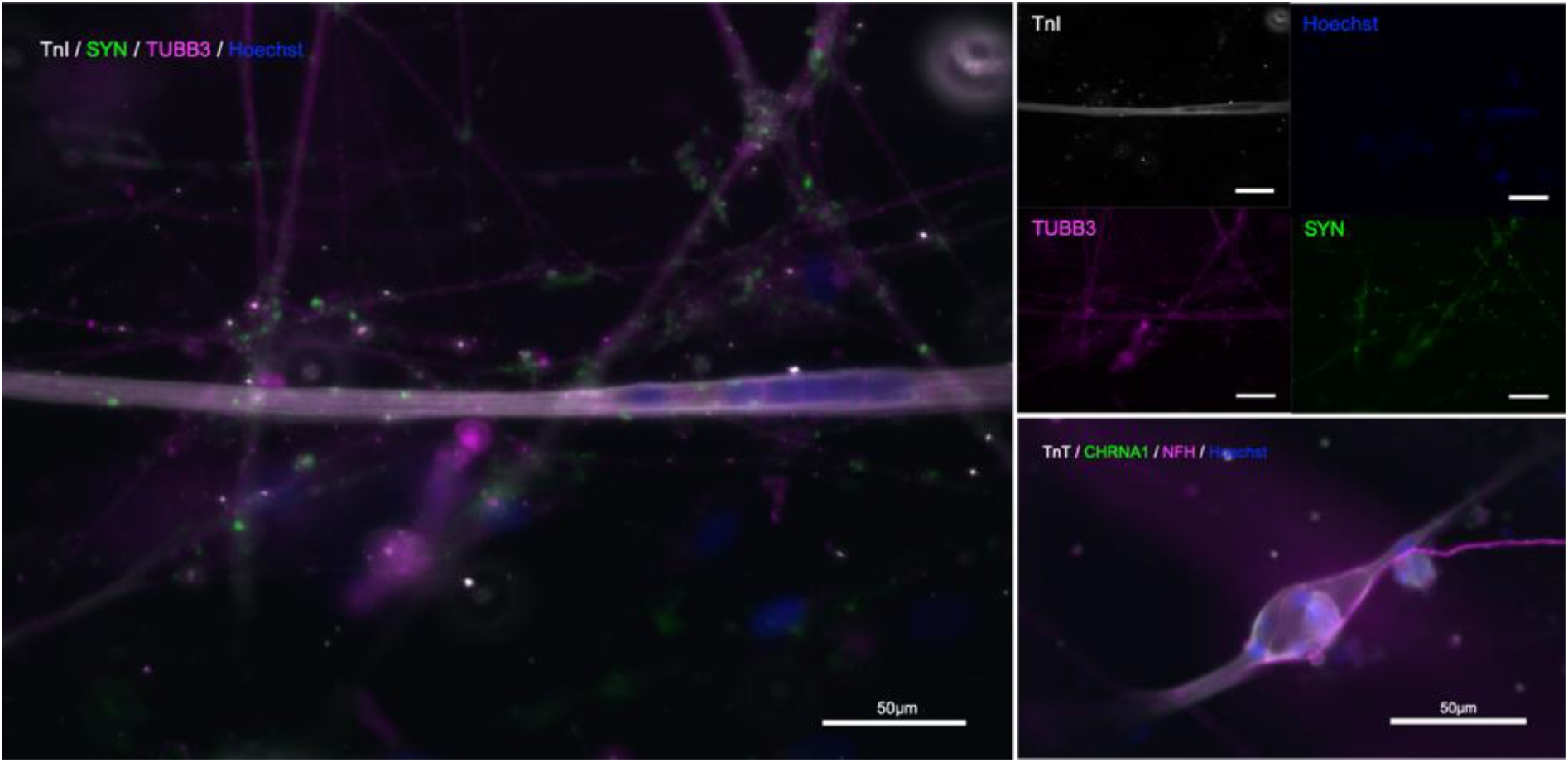
Immunostaining in microfluidic chips indicates presence of NMJs. (**a**) and (**b**): Troponin I (TnI, grey), synaptophysin (SYP, green), and β-III-tubulin (TUBB3, magenta), nuclear stain Hoechst (blue). (**c**) neurofilament-heavy (NF-H, magenta), nicotinic acetylcholine receptor subunit 1 (CHRNA1, green), troponin T (TnT, grey), nuclear stain Hoechst (blue).

**Figure 6.**
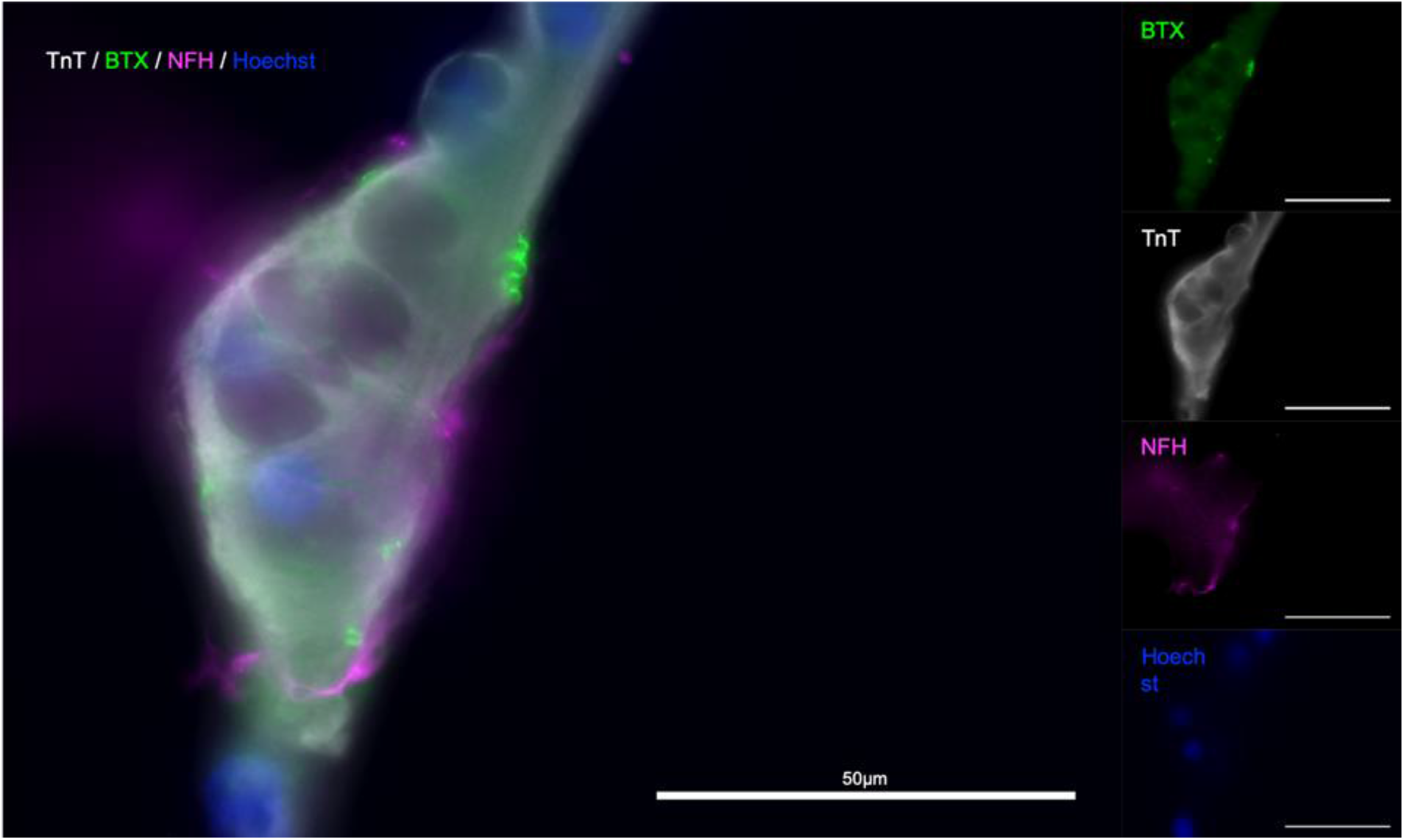
α-Bungarotoxin staining indicates presence of functional NMJs. Troponin T (TnT, grey), α-Bungarotoxin (BTX, green), and Neurofilament-heavy (NF-H, magenta), nuclear stain Hoechst (blue). Merged

### Alpha-Bungarotoxin abolishes observable contractile activity in the *in vitro* NMJs

To confirm that the contractile activity observed was due to the presence of functional NMJs, we made use of the natural functionality of α-Bungarotoxin which causes muscle paralysis by blocking activity of the postsynaptic nicotinic acetylcholine receptors in the NMJ. We next wanted to see whether the contractions we were observing in the myotube well could be inhibited by introducing α-Bungarotoxin to the cultures. In a pilot experiment, we observed a complete abolition of contractile activity in all experimental group chips, while the control saw a slight increase in contraction frequency. In a blinded follow-up, similar responses were observed within 3 of the myotubes imaged in the experimental group (n=6) having their activity completely abolished, one with a reduction from 58 to 1 contraction in the first minute of the observed window, one with a reduction of contractile frequency by over 91% (124 contractions per 10 min to 11), and the last with short bursts of increased contraction frequency that occurred in groups with an increasing proportion of partial contractions and contractile activity ceasing completely after 6.5 min. Similarly, the myotube that showed a 91% decrease in activity performed one full contraction and 10 partial contractions thereafter (Figure 7). Subsequent imaging of the chips revealed the presence α-Bungarotoxin binding on Tn-positive cells that were previously observed to lose their contractile activity in the experiment (Figure 8), in close proximity to axons (beta-3-tubulin staining).

**Figure 7.**
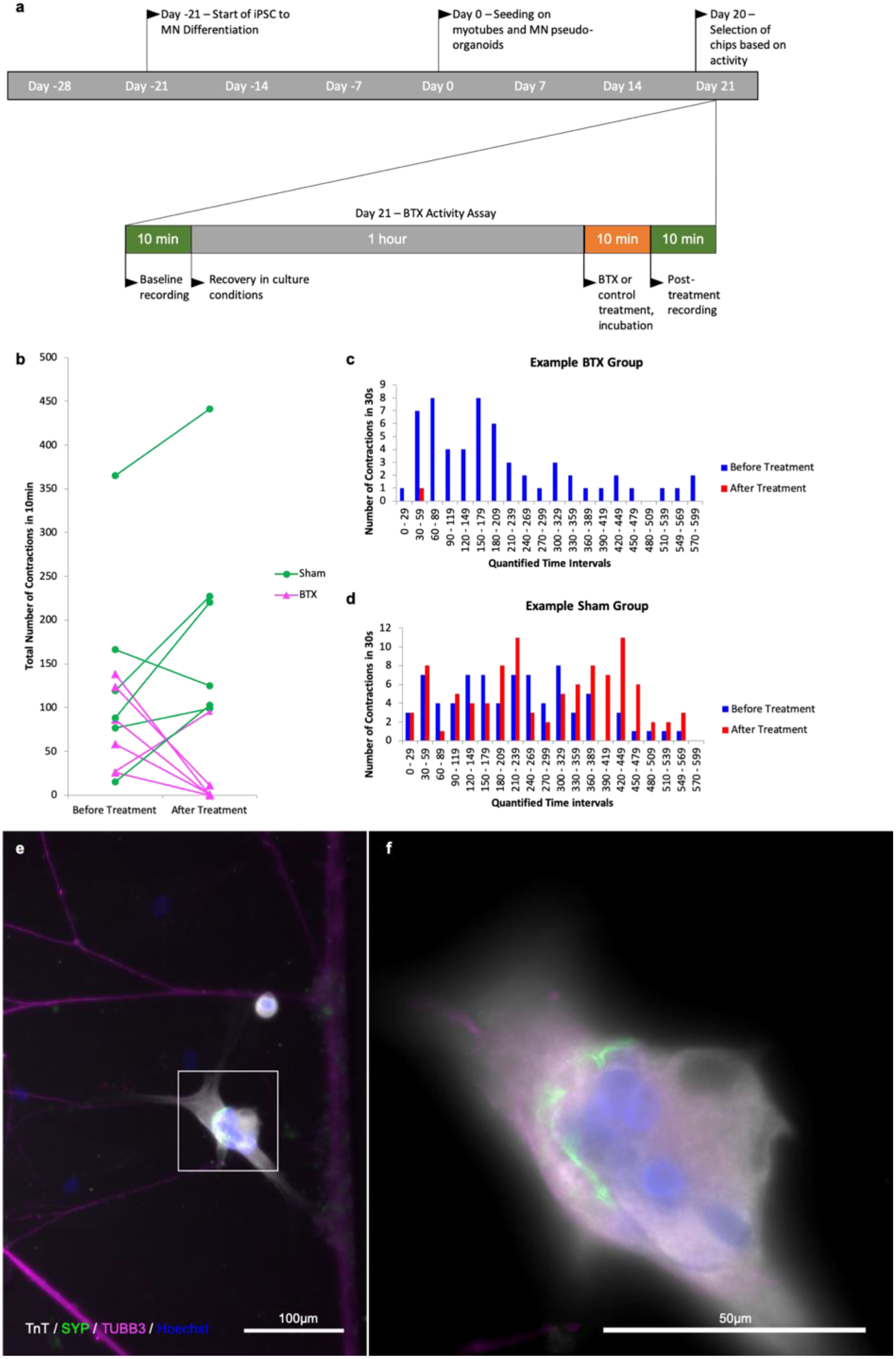
Quantification of contractile activity before and after α-Bungarotoxin application BTX binding at myotube surface. Outline of experimental timeline (**a**), total number of contractions in a 10 min time interval in a myotube in contact with one or more axons in the control and BTX-treated groups before and after treatment (**b**) number of observed myotube contractions in 30s intervals before and after treatment in a myotube in the BTX-treated (**c**) and sham-treated (**d**) groups. (**e**) Image of myotube and MN axons after BTX application. (**f**) Higher magnification of the myotube in the area boxed in (**e**); Troponin T (TnT, grays), α-Bungarotoxin (BTX, green), and beta-III-tubulin (magenta), nuclear stain Hoechst (blue).

**Figure 8.**
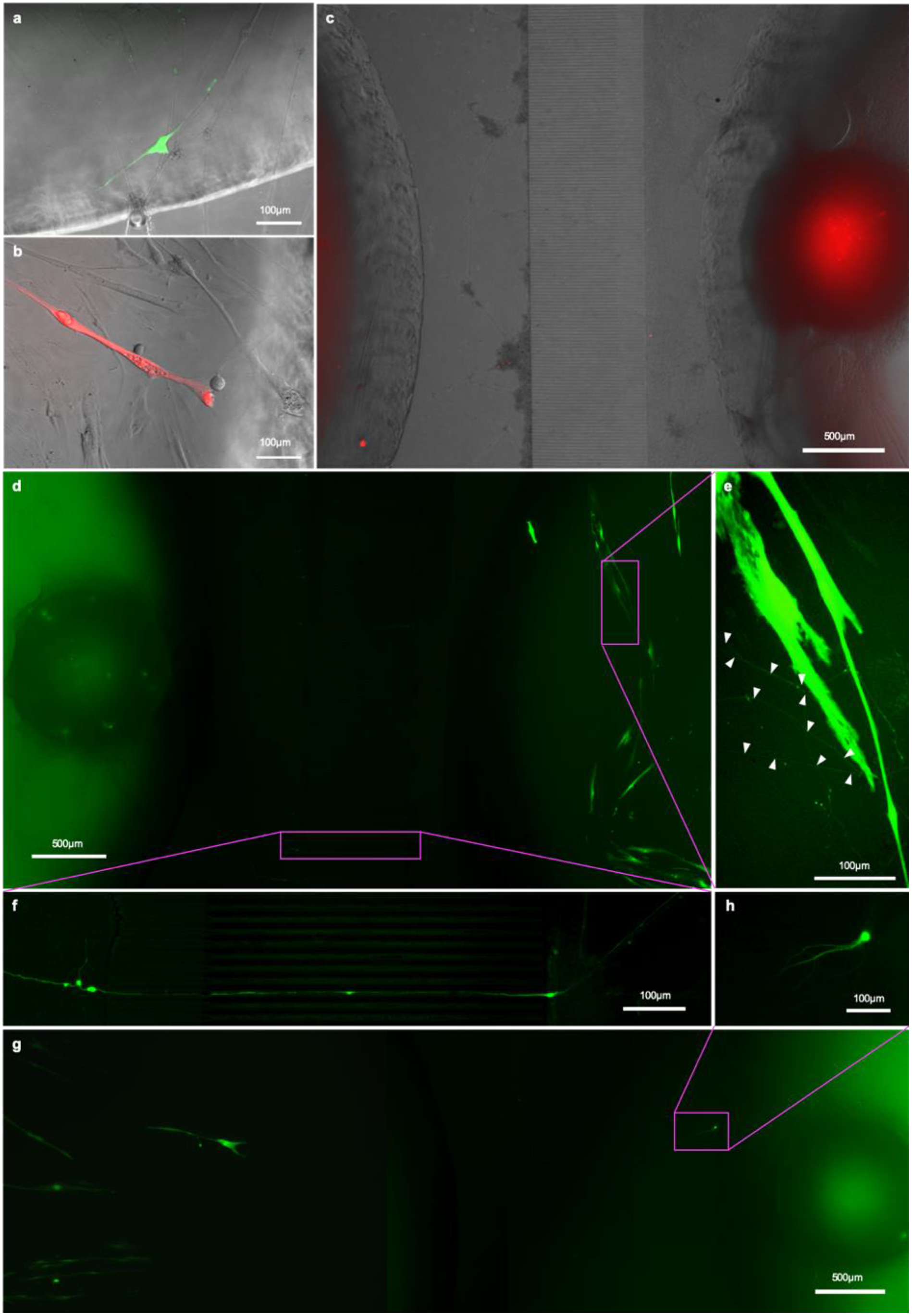
RV-infection and retrograde tracing. (**a**) EnvA-ΔG-RV-GFP and (**b**) EnvA-ΔG-RV-mCherry-infected myotube fluorescence 48h after infection superimposed on phase-contrast images of the respective myotubes; (**c**, **d**, **g**) overview tracing EnvA-ΔG-RV-mCherry fluorescence superimposed on phase-contrast image at 31 DIV (**c**) and EnvA-ΔG-RV-GFP (**d**, **g**) from infected myotubes to cell bodies inside the aggregates (**d**) and cells that migrated out (**g**, **h**) at 29 DIV; (**e**) details of axons connecting up the infected myotubes and (**f**) crossing the channels from the outer to the myotube well.

### Retrograde tracing with pseudotyped rabies virus shows presence of functional NMJs

Finally, we wanted to see whether we could demonstrate NMJ functionality by exploiting the natural infection pathway of rabies from muscle to MNs, employing fluorescently labelled, glycoprotein-deleted, pseudotyped rabies virus for monosynaptic retrograde tracing (Wickersham et al, 2010, Osakada and Callaway, 2013).

To this end, we transfected the myoblasts in the central well of our chips with helper constructs pCAG-YTB^24^ or pAAV-CMV-TVAmCherry-2A-oG^25^ by electroporation to express the exogenous receptor TVA and rabies-G immediately before seeding them in the well. Otherwise the chips were seeded as before and the cultures allowed to mature. After 15DIC the cells in the myotube well were infected with either EnvA-ΔG-RV-mCherry or EnvA-ΔG-RV-GFP (for pCAG-YTB or pAAV-CMV-TVAmCherry-2A-oG groups, respectively). The rationale for this is that myotubes expressing the helper construct are infected by EnvA-ΔG-RV and subsequently express the trasngenes and form virions. Only MNs connected to rabies infected myotubes will receive virions through the synapses. In both experimental groups, the first myotubes showed bright fluorescence by 17 DIV (Figure 9 A-B) however, not all transfected myotubes were infected and the myotubes that were infected started becoming less healthy and dying after a few days. By 20 DIC, the first neurons in the pAAV-CMV-TVAmCherry-2A-oG and EnvA-ΔG-RV-GFP group showed observable levels of fluorescence. From 20 DIV until the experiment was terminated at 31 DIV, we observed one or more neurons in at least one pseudo-organoid in each of the pAAV-CMV-TVAmCherry-2A-oG chips becoming infected, with the infected neurons typically dying within a week of becoming bright enough to detect. Somata of the infected neurons were found both amongst the cells that had migrated out from the aggregate (Figure 9 H) as well as throughout the aggregate itself (Figure 9 D). In the pCAG-YTB group we observed infection of MNs in 2/5 chips, with the first detectable infection only occurring 10 days after initial infection of the myotube well.

## Discussion and Conclusions

In this study, we demonstrate that we can reproducibly generate functional NMJs from human iPS cell-derived MNs and primary myotubes using an in-house developed microfluidic chip. The specific microfluidic platform offers a particular advantage to other previously described approaches in that the small axon channels with controllable connectivity allow for near complete isolation of extracellular environments through use of hydrostatic pressure, thus enabling chemical insult to NMJs and MN cell bodies to be carried out independently.^13,14^ Furthermore, to the best of the authors’ knowledge, this is the first study demonstrating validation of *in vitro* NMJ functionality through monosynaptic retrograde tracing with a designer glycoprotein-deleted rabies virus, employing two different helper constructs to allow virus entry to the myotubes and producing infective virus particles. Monosynaptic retrograde tracing, correlated with the abolition of contractile activity in the NMJ using α-BTX, provides unequivocal evidence of the functionality of the NMJs engineered within the microfluid chip.

Having verified the antigenic profile of iPS cell-derived MNs, we proceeded to establish their functionality by assessing the spontaneous activity of MN networks on MEAs over time. Key parameters such as firing rate across the entire MEA, together with BI, provide a good overall impression of the state of the network, with regard to emergence of spontaneous electrical activity and synchronization, respectively.^26^ Identification of three distinct stages of MN network maturity suggests that early increases in overall activity in the first five weeks does not indicate the establishment of a well-connected network, as significant bursting activity is not observed until 42DIV, at which point the firing rate is already quite high and does not significantly increase further. Following this, even though overall activity decreases, as seen in the firing rate, the activity is highly synchronised and mostly contained in burst firing, consistent with previous findings.^26^ Taken together, these results are important as they show that the iPS cell-derived MNs form functional, coordinated networks before proceeding with the *in vitro* NMJ assays.

To unequivocally establish the functionality of the in vitro NMJs, we chose to deliver the TVA and rabies glycoprotein-containing single helper constructs, required to allow infection of the myoblasts by the pseudotyped ΔG-rabies virus and production of infectious viral particles, respectively, by electroporation over other delivery methods. The myotubes expressing the helper construct will be infected on application of EnvA-ΔG-RV, subsequently express transgenes and form virions. Only MNs synaptically connected with the rabies expressing myotubes will receive virions through the synaptic contact between MNs and infected myotubes. Virus expressing MNs cannot form infectious virus particles due to lack of rabies-G, hence the viruses will not propagate further from one infected MN to another. This approach avoids any risk of chance infection of one of the axons extending into the myotube chamber of our microfluidic chips. The lower infection efficiency in the group using pCAG-YTB as a vector for myotube infection was not unexpected as the helper construct is an older version than the recently optimized pAAV-CMV-TVAmCherry-2A-oG. Notwithstanding this issue and the fact that the use of electroporation may not be optimal for construct delivery, we still managed to see transmission of the virus from the infected myotubes to MNs with both constructs. This supports the notion that our model provides a robust platform for investigating mechanisms involving NMJ formation and function. It is noteworthy that, while there were some infected MNs that had migrated out of the MN aggregates (presumably after having established contact with the myotubes), most of the infected cells were located inside the pseudo-organoids. Interestingly, some of the cell bodies that showed fluorescence were located on the far side of the MN aggregates making visual detection challenging.

It was repeatedly observed that RV-infected myotubes contacted by one or more axons would become brightly fluorescent, detach and die in the days before the associated axons and respective cell bodies became fluorescent enough to visualize and trace. Given the brightness of the fluorescence usually observed before this occurred, it was not unexpected, as it correlates with the extent to which the cellular machinery is taken over by the virus for production of more viral particles and the corresponding load on homeostatic mechanisms.^25^

α-BTX has been shown to have a similarly high affinity to the neuronal nACHRs subunits α7 and α9/10.^27^ The ease of controlling for this in our platform, through ensuring unidirectional flow of media from the outer to the inner well through hydrostatic pressure, demonstrates the practical advantage of this feature of the microfluidic chip and emphasises the possibilities for employing the overall model in selectively inducing perturbations that mimic aspects of NMJ dysfunction as part of ALS pathology. This is a promising perspective in the context of developing robust preclinical models that may help address some of the limitations of *in vivo* ones.

The addition of α-BTX completely abolished contractile activity in the *in vitro* NMJs, except in one the units, in which some contractile activity could be observed. The latter was not entirely surprising, given that cultured myotubes tend to show spontaneous activity in culture.^28,29^ It was particularly of note that the kind of contraction observed changed in the myotubes that did not show complete abolishment of activity. In one case the activity was reduced to one full contraction and ten partial contractions in two groups as opposed to a mix of both occurring throughout the observed period before addition of α-BTX. This incomplete or delayed effect may have been due to a delay in mixing speed between bulk media in the well and the narrow space within the active zone of the microfluidic chip, which, in some cases may have been exacerbated by the presence of large axon bundles around the myotube.

Preclinical *in vivo* models largely involve the generation of transgenic mice overexpressing some of the most common gene mutations associated with ALS. For example, various transgenic mutant mouse lines overexpressing the SOD1 mutation have been generated. These mutants recapitulate hallmarks of ALS pathology, such as limb tremor, locomotor deficits, and paralysis.^3^ Similarly, the TDP-43 mutant mouse models display positive cytoplasmic inclusions of the protein^5^, while FUS models manifest progressive, mutant-dependent MN degeneration^7^ and functional abnormalities in the NMJ^6^. On the other hand, while C9ORF72 mutants may display evidence of ALS-related pathology, such as a microglia phenotype or frontotemporal dementia (FTD), they fail to recapitulate MN degeneration.^7^ Thus, while the SOD1, TDP-43, FUS, and C90RF72 mouse mutant lines may afford highly useful insights into specific pathological features and disease mechanisms of ALS, no single model can fully capture the immense complexity of the disease or faithfully mimic the human patient organism. Similar limitations apply to other vertebrate, as well as invertebrate models of ALS. ^30–32^

In light of the above, *in vitro* models, including NMJ engineering approaches are highly relevant for studying and elucidating mechanistic causes of the disease. The differentially-perturbable microfluidic environment applied in our study thus represents a highly versatile platform for *in vitro* modelling of ALS pathology, also in the context of patient-specific modelling. Employing the platform in combination with an integrated MEA interface should further increase its usefulness by allowing for electrophysiological analysis of the system as well as its responses to selective perturbations. Additionally, these platforms can be applied for preliminary drug screening. As such, the platforms are highly complementary to *in vivo* disease modelling, while they also have the potential of significantly reducing or replacing animal models.

## Supporting information

HB9 and ISL1 expression in Sections of MN aggregates

## References

1. Johnston, C. A. et al. Amyotrophic lateral sclerosis in an urban setting: A population based study of inner city London [2]. J. Neurol. 253, 1642–1643 (2006).

2. Hirtz, D. et al. How common are the “common” neurologic disorders? Neurology 68, 326 LP–337 (2007).

3. Nardo, G. et al. Immune response in peripheral axons delays disease progression in SOD1 G93A mice. J. Neuroinflammation 13, 1–16 (2016).

4. De Vos, K. J. & Hafezparast, M. Neurobiology of axonal transport defects in motor neuron diseases: Opportunities for translational research? Neurobiol. Dis. 105, 283–299 (2017).

5. Kabashi, E. et al. TARDBP mutations in individuals with sporadic and familial amyotrophic lateral sclerosis. Nat. Genet. 40, 572–574 (2008).

6. Sharma, A. et al. ALS-associated mutant FUS induces selective motor neuron degeneration through toxic gain of function. Nat. Commun. 7, 1–14 (2016).

7. Philips, T. & Rothstein, J. D. Rodent Models of Amyotrophic Lateral Sclerosis. Curr. Protoc. Pharmacol. 69, 5.67.1–5.67.21 (2015).

8. O’Rourke, J. G. et al. C9orf72 BAC Transgenic Mice Display Typical Pathologic Features of ALS/FTD. Neuron 88, 892–901 (2015).

9. Takahashi, K. & Yamanaka, S. Induction of Pluripotent Stem Cells from Mouse Embryonic and Adult Fibroblast Cultures by Defined Factors. Cell 126, 663–676 (2006).

10. Abernathy, D. G. et al. MicroRNAs Induce a Permissive Chromatin Environment that Enables Neuronal Subtype-Specific Reprogramming of Adult Human Fibroblasts. Cell Stem Cell 21, 332–348.e9 (2017).

11. Richner, M., Victor, M. B., Liu, Y., Abernathy, D. & Yoo, A. S. MicroRNA-based conversion of human fibroblasts into striatal medium spiny neurons. Nat. Protoc. 10, 1543–1555 (2015).

12. Amoroso, M. W. et al. Accelerated high-yield generation of limb-innerv. J. Neurosci. 33, 574–586 (2013).

13. van de Wijdeven, R. et al. Structuring a multi-nodal neural network in vitro within a novel design microfluidic chip. Biomed. Microdevices 20, (2018).

14. van de Wijdeven, R. et al. A novel lab-on-chip platform enabling axotomy and neuromodulation in a multi-nodal network. Biosens. Bioelectron. 140, 111329 (2019).

15. Park, H. S., Liu, S., McDonald, J., Thakor, N. & Yang, I. H. Neuromuscular junction in a microfluidic device. Proc. Annu. Int. Conf. IEEE Eng. Med. Biol. Soc. EMBS 2833–2835 (2013). doi:10.1109/EMBC.2013.6610130

16. Morimoto, Y., Kato-Negishi, M., Onoe, H. & Takeuchi, S. Three-dimensional neuron-muscle constructs with neuromuscular junctions. Biomaterials 34, 9413–9419 (2013).

17. Demestre, M. et al. Formation and characterisation of neuromuscular junctions between hiPSC derived motoneurons and myotubes. Stem Cell Res. 15, 328–336 (2015).

18. Ionescu, A., Zahavi, E. E., Gradus, T., Ben-Yaakov, K. & Perlson, E. Compartmental microfluidic system for studying muscle-neuron communication and neuromuscular junction maintenance. Eur. J. Cell Biol. 95, 69–88 (2016).

19. Happe, C. L., Tenerelli, K. P., Gromova, A. K., Kolb, F. & Engler, A. J. Mechanically patterned neuromuscular junctions-in-a-dish have improved functional maturation. Mol. Biol. Cell 28, 1950–1958 (2017).

20. Lewandowska, M. K., Bogatikov, E., Hierlemann, A. R. & Punga, A. R. Long-term high-density extracellular recordings enable studies of muscle cell physiology. Front. Physiol. 9, 1–12 (2018).

21. Mills, R. J. et al. Development of a human skeletal micro muscle platform with pacing capabilities. Biomaterials 198, 217–227 (2019).

22. Wickersham, I. R., Sullivan, H. A. & Seung, H. S. Production of glycoprotein-deleted rabies viruses for monosynaptic tracing and high-level gene expression in neurons. Nat. Protoc. 5, 595 (2010).

23. Osakada, F. & Callaway, E. M. Design and generation of recombinant rabies virus vectors. Nat. Protoc. 8, 1583 (2013).

24. Marshel, J. H., Mori, T., Nielsen, K. J. & Callaway, E. M. Targeting single neuronal networks for gene expression and cell labeling in vivo. Neuron 67, 562–574 (2010).

25. Ciabatti, E., González-Rueda, A., Mariotti, L., Morgese, F. & Tripodi, M. Life-Long Genetic and Functional Access to Neural Circuits Using Self-Inactivating Rabies Virus. Cell 170, 382–392.e14 (2017).

26. Wagenaar, D. A. Controlling Bursting in Cortical Cultures with Closed-Loop Multi-Electrode Stimulation. J. Neurosci. 25, 680–688 (2005).

27. Couturier, S. et al. A neuronal nicotinic acetylcholine receptor subunit (α7) is developmentally regulated and forms a homo-oligomeric channel blocked by α-BTX. Neuron 5, 847–856 (1990).

28. Guo, X. et al. In vitro differentiation of functional human skeletal myotubes in a defined system. Biomater. Sci. 2, 131–138 (2014).

29. Al Samid, M. A. et al. A functional human motor unit platform engineered from human embryonic stem cells and immortalized skeletal myoblasts. Stem Cells Cloning Adv. Appl. 11, 85–93 (2018).

30. Matthaei, K. I. Genetically manipulated mice: A powerful tool with unsuspected caveats. J. Physiol. 582, 481–488 (2007).

31. Watson, M. R., Lagow, R. D., Xu, K., Zhang, B. & Bonini, N. M. A Drosophila model for amyotrophic lateral sclerosis reveals motor neuron damage by human SOD1. J. Biol. Chem. 283, 24972–24981 (2008).

32. Perrin, S. Make mouse studies work. Nature 507, 423–425 (2014).

