## Supplementary material for "Modelling functional human neuromuscular junctions in a differentially-perturbable microfluidic environment, validated through recombinant monosynaptic pseudotyped ΔG-rabies virus tracing": HB9 and ISL1 expression in Sections of MN aggregates

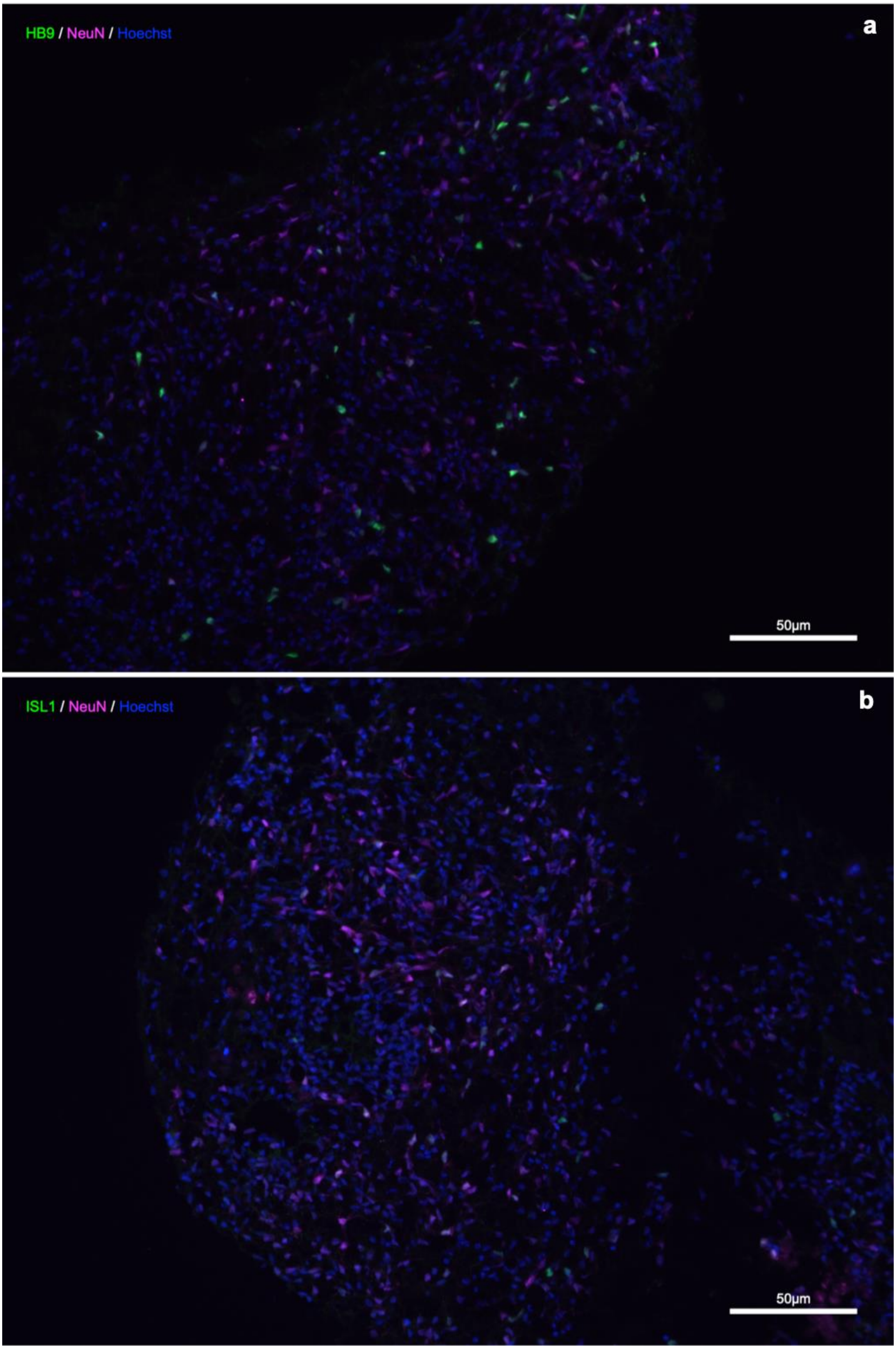
HB9 / NeuN / Hoechst
a
50µm
ISL1 / NeuN / Hoechst
b
50µm

**Supplementary Figure 1. HB9 and ISL1 expression in Sections of MN aggregates.**
Sections of MN aggregates immunolabelled for NeuN (magenta) and (**a**) Homeobox HB9 (HB9, green) or Islet 1 (ISL1, green), nuclear staining with Hoechst (Blue)
